# Transcriptomic analyses reveal neuronal specificity of Leigh syndrome associated genes

**DOI:** 10.1101/2022.08.05.502943

**Authors:** Azizia Wahedi, Chandika Soondram, Alan E. Murphy, Nathan Skene, Shamima Rahman

## Abstract

Leigh syndrome (subacute necrotising encephalomyelopathy) is a rare inherited, complex, and typically early onset mitochondrial disorder with clinical and genetic heterogeneity. It owes its heterogeneous nature to the complex nature of mitochondrial genetics and the significant interactions that occur between the mitochondrial and nuclear genomes. Stepwise developmental regression is a classical feature of patients with Leigh syndrome, but other neurological features include ataxia, tremor, seizures, and occasionally psychiatric manifestations. To date the involvement of more than 100 genes has been identified in patients with Leigh syndrome. Despite this, the pathophysiology of Leigh syndrome remains unknown, although it is thought to be due to primary mitochondrial dysfunction.

Here, we sought to determine the cellular specificity of all Leigh syndrome associated genes within the brain and investigate a potential common genetic mechanism between Leigh syndrome and other disorders with overlapping clinical features. We utilized co-expression network analyses constructed using existing UK Brain Expression Consortium (UKBEC) data from 10 brain regions to identify regions of the brain where our genes of interest were enriched. Next, expression weighted cell type enrichment (EWCE) was employed to determine the cellular specificity of Leigh syndrome associated genes. Association with pre-or post-synaptic structures was identified using synaptic gene ontology (SynGO) analysis. Finally, heritability analysis was performed on the co-expression network modules demonstrating enrichment of Leigh syndrome associated genes to investigate potential genetic relationships with Parkinson disease, epilepsy, and schizophrenia.

Co-expression network analyses reveal that genes encoding oxidative phosphorylation subunits and assembly factors exhibit the highest levels of expression in brain regions most affected on MRI and neuropathology in Leigh syndrome. Further, over two thirds of the genes were found to be enriched within the substantia nigra and 19 genes appeared to cluster, with selective enrichment in the putamen, substantia nigra, medulla and thalamus. EWCE analyses of single cell RNA-Seq data from mouse brain revealed significant enrichment of Leigh syndrome associated genes in hippocampal and somatosensory pyramidal neurones and interneurons of the brain. SynGO analysis demonstrated that expression of these genes is preferentially associated with pre-synaptic structures. Heritability studies suggested that some modules in which Leigh syndrome associated genes were enriched were also enriched for Parkinson disease and epilepsy.

In conclusion, our findings suggest a primary mitochondrial dysfunction as the underlying basis of Leigh syndrome with causative genes primarily expressed in neuronal cells, explaining the relative sparing of glial cells observed in neuropathological studies.

## Introduction

Leigh syndrome (subacute necrotizing encephalomyelopathy), first described in 1951, is a severe, early-onset inherited neurometabolic disorder which primarily affects the central nervous system. With a minimum birth prevalence of 1 in every 40,000 live births^1,2^, Leigh syndrome is the most frequent clinical presentation of mitochondrial disease in childhood. Onset typically varies between 3 and 12 months of age, before ultimately leading to early death^3^. Survival rates vary according to the specific gene defect, but one large multinational study reported a median age of death of approximately 2.4 years across the whole cohort^3^. Patients affected by Leigh syndrome typically experience global developmental delay and/or regression, coupled with seizures, ataxia, hypotonia, dystonia, and optic atrophy. These neurological symptoms are related to progressive bilateral, often symmetrical lesions affecting the basal ganglia, midbrain, and brainstem^1,4^. The cerebellum, white matter and spinal cord may also be affected in some cases. In addition, Leigh syndrome was associated with multisystemic disease manifestations in some patients who presented hepatic, renal, and cardiac symptoms.

Since the advent of next-generation exome and genome sequencing, an increasing number of genetic causes of Leigh syndrome has been identified^5^. As of 2021, a total of 113 disease-causing genes have been reported (https://search.clinicalgenome.org/kb/conditions/MONDO:0009723) reflecting the phenotypic and biochemical heterogeneity of the disorder. The genetic causes encompass mutations in both mitochondrial DNA (mtDNA), which may be maternally inherited or sporadic, and nuclear DNA, which are usually inherited in an autosomal recessive pattern (but may rarely be dominant or X-linked). As far as mtDNA mutations are concerned, they can either be homoplasmic or heteroplasmic (i.e., co-existence of mutated and wild-type mtDNA), and the mutation load appears to, at least partly, determine the phenotype. Most patients with Leigh syndrome have causative mutations in nuclear DNA genes, whilst mtDNA mutations are responsible in approaching 25% of cases^3^.

Leigh syndrome and other mitochondrial disorders form a group of approaching 400 different rare monogenic disorders unified by impacts on mitochondrial structure and function^6^. These mitochondrial disorders are characterised by extreme clinical heterogeneity, leading to considerable diagnostic odysseys for affected patients. Moreover, there is currently a dearth of effective disease-modifying therapies. The aim of this study was to investigate the cellular specificity of Leigh syndrome associated genes within the brain, which may ultimately facilitate the identification of novel therapeutic targets.

This project involved using a systems biology approach to explore the cellular specificity of 113 genes linked to Leigh syndrome, which have recently been curated by a global ClinGen expert panel (https://search.clinicalgenome.org/kb/conditions/MONDO:0009723). In summary, a transcriptomic data set was utilised used to (i) compare gene expression levels across 10 UKBEC brain regions available in transcriptomic data from the United Kingdom Brain Expression Consortium (UKBEC) network, (ii) investigate the cell type(s) which are most relevant to the disease, (iii) identify whether the genes enriched in selected modules are associated with presynaptic or postsynaptic structures, and (iv) test for genetic commonalities, if any, between Leigh syndrome and other neurological disorders.

## Materials and methods

### Co-expression network analysis

CoExp is a resource developed to exploit gene co-expression networks (GCNs) to build gene groupings based on transcriptomic profiling^7^. The CoExp tool is an online web application consisting of a collection of 109 networks, powered by the CoExpNets suite of R packages. The enrichment of the 113 genes of interest (Table 1) was tested within the co-expression networks generated from different brain regions, available in the CoExp Web Application (https://rytenlab.com/coexp). The ten human brain regions within the UKBEC network which were investigated were as follows: cerebellar cortex (CRBL), frontal cortex (FCTX), hippocampus (HIPP), medulla oblongata (MEDU), occipital cortex (OCTX), putamen (PUTM), substantia nigra (SNIG), temporal cortex (TCTX), thalamus (THAL), and white matter (WHMT). The UKBEC data set comprises 134 individuals confirmed to be neuropathologically normal and of European descent from 16 to 102 years of age and was generated using Affymetrix Exon 1.0 ST Arrays^8^.

The Gene Annotation functionality of the CoExp website returns a table with detailed information regarding the genes present and the modules in which they are found within a specific network. The FDR-adjusted *P*-value was used for analysis, as described by Mencacci et al.^9^, where values < 0.05 satisfied the argument that those genes were significantly enriched within the specific modules. For 79 genes of interest that were available in the UKBEC network, specific modules in which Leigh syndrome associated genes were significantly enriched were identified (Table 2).

### Venny analysis

A Venny diagram (https://bioinfogp.cnb.csic.es/tools/venny/) was used to test for overlapping of Leigh syndrome genes amongst enriched modules and to summarise the CoExp findings from Table 3. Only the genes within modules with enrichment of the largest number of Leigh associated genes (substantia nigra, putamen, medulla, and thalamus) were used as input for the diagram.

### Expression-weighted cell-type enrichment analysis

Expression-weighted cell-type enrichment (EWCE) analysis was performed to investigate whether certain cell types express the Leigh syndrome associated genes more than can be attributed to random chance^10^. In this study, we used the EWCE method with the target gene list (Table 1) and background gene set (defined as all other genes within the data set) as the two arguments to determine the probability that the 113 genes exhibit higher specificity in a given cell type. The data set within the EWCE package includes the Karolinska Institute single cell RNA-Seq data from mouse (juvenile P22-P32 CD1 *Mus musculus*) brain^11^, thus genes in our target list were converted to their mouse homologs, thorough a function within EWCE that converts the HGNC gene name into its corresponding MGI mouse ortholog gene. Based on the standard deviation from the mean, the enrichment of Leigh syndrome genes in level 1 and level 2 cell types was determined, with the levels representing higher (cell types) and lower level (cell subtypes) groupings, respectively. To investigate whether subgrouping the genes based on product function would reveal differential enrichment, we re-ran the EWCE analysis on functional groupings of the Leigh syndrome associated genes (Table 1). We were unable to perform the same analysis with the genes subdivided based on whether the mode of inheritance is mitochondrial or nuclear because no mitochondrial encoded genes are present within the EWCE dataset. Instead, we utilised the online STRING database platform to identify predicted direct (physical) and indirect (functional) associations of the products of the protein coding mitochondrial genes linked to Leigh syndrome. The corresponding genes for proteins interacting with our target gene list products were compiled and utilised as an extended ‘mitochondrial’ gene set and then run through EWCE. EWCE was run on 10,000 iterations for the complete list of genes and was increased to 100,000 bootstrap replicates to compensate for the multiple testing. We also controlled for transcript length and GC content of the genes. Data are displayed as standard deviations from the mean and p-values were obtained using the Benjamini-Hochberg FDR method to account for multiple testing.

Next, we investigated three cell populations in which Leigh syndrome associated genes were found to be significantly enriched, namely, pyramidal CA1, pyramidal SS, and interneurons, to determine the specificity values of the genes. Specificity is defined as the average expression of a gene in a particular cell type by the average expression of the same gene in all other cell types. The specificity values of Leigh syndrome genes in the respective cell types were obtained from a previous publication^12^.

### Synaptic association

To investigate the association of Leigh syndrome enriched modules with presynaptic and/or postsynaptic structures, the modules listed in Table 2 were annotated in the CoExp website and then inputted in the Synaptic Gene Ontologies (SynGO)^13^ website (https://syngoportal.org/). The latter analysed the gene list for synaptic association, compared to a default background of all brain expressed genes (GTExV7 brain tissue), using a one-sided Fisher exact test. Once a term particularly enriched with the gene cluster was found, multiple testing corrections were applied using FDR and the resulting plots were coloured according to the enrichment q-values.

### Heritability enrichment analysis

A heritability study was performed to assess the potential co-morbidity of Leigh syndrome with neurological disorders characterised by overlapping clinical features (such as dystonia and seizures). Candidate genes for Parkinson disease^14^, epilepsy^15^, and schizophrenia^16^ were obtained from the literature and FDR-adjusted P-values were derived from the CoExp Gene Set Annotation tool. The co-expression modules in Table 2, containing more than 15 Leigh syndrome associated genes, were selected and the FDR-adjusted *P*-values for the different neurological disorders were used for the plot, with a nominal enrichment threshold of *P*<0.05.

### Data availability and URLs

Data used to generate Figures 1, 2, and 5 are available within the Supplementary materials (Supplementary Tables 1, 2, and 4, respectively). The level 1 (cell type) and level 2 (cell subtypes) specificity values for the Karolinska single-cell mouse RNA-Seq superset can be retrieved online from Nathan Skene’s GitHub (https://github.com/NathanSkene/MAGMA_Celltyping). All gene groupings, enrichment annotations and heritability analyses were performed using the CoExp website (www.rytenlab.com/coexp/Run/Catalog/), along with its supplementary CoExpNets packages (https://github.com/juanbot/CoExpNets). Gene lists from the CoExp tool were inputted into the Venny 2.1 online tool (https://bioinfogp.cnb.csic.es/tools/venny/) and the freely accessible SynGO website (https://www.syngoportal.org/) for additional analyses.

**Figure 1.**
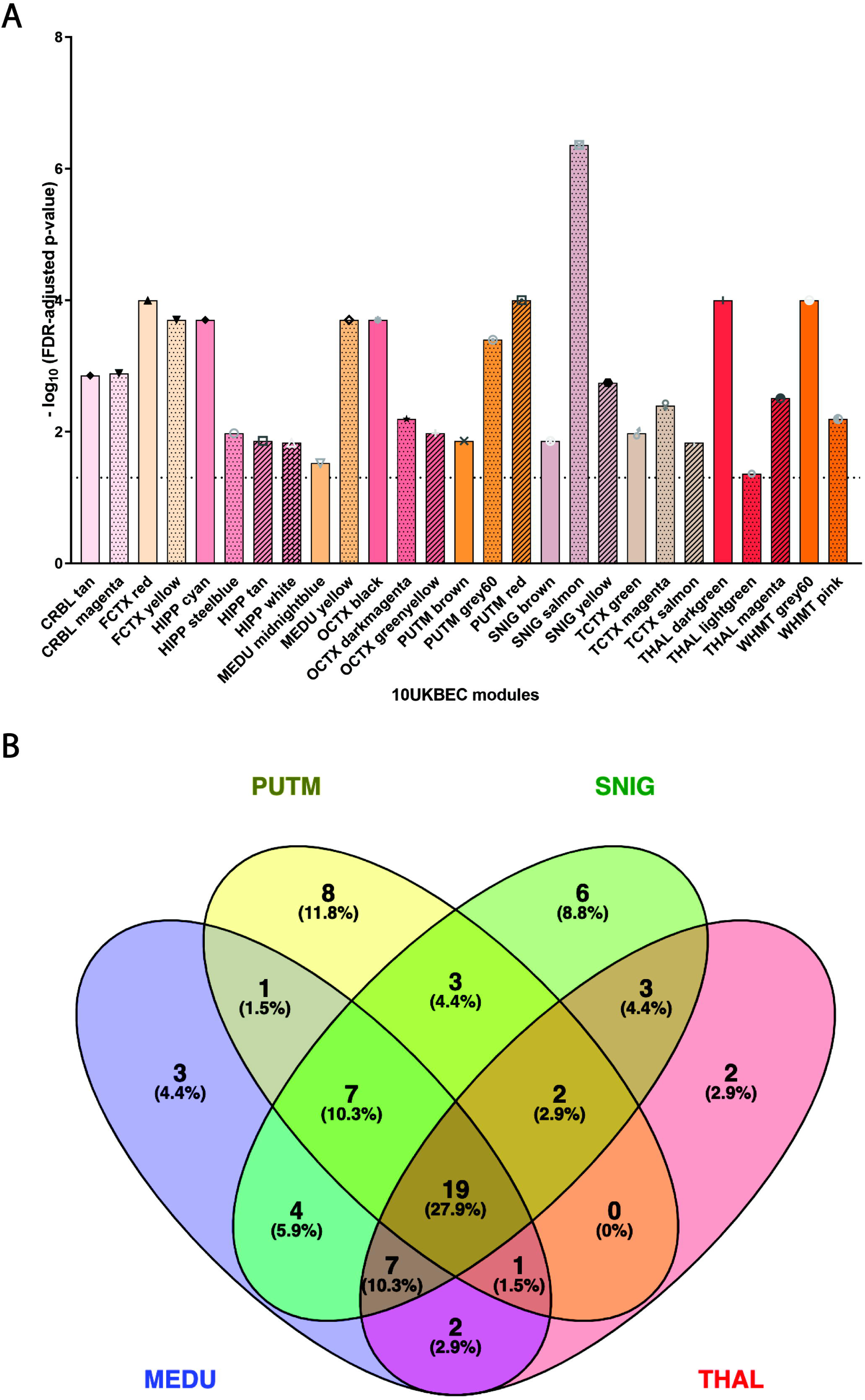
Expression of Leigh syndrome associated genes within the enriched UKBEC modules. **(A)** Expression of Leigh syndrome associated genes in UKBEC network co-expression modules **(B)** Venny diagram showing that 64 % of the genes within four regions of interest are enriched within the substantia nigra with one or more overlapping modules, and almost 10% of the genes are only expressed in the substantia nigra. This suggests that the substantia nigra is likely to be the most affected in patients with Leigh syndrome owing to its high number of Leigh syndrome-associated genes. The dashed line marks the approximate FDR threshold cut-off for nominal enrichment (approximately 1.301). Key: MEDU = medulla, PUTM = putamen, SNIG = substantia nigra, THAL = thalamus

**Figure 2.**
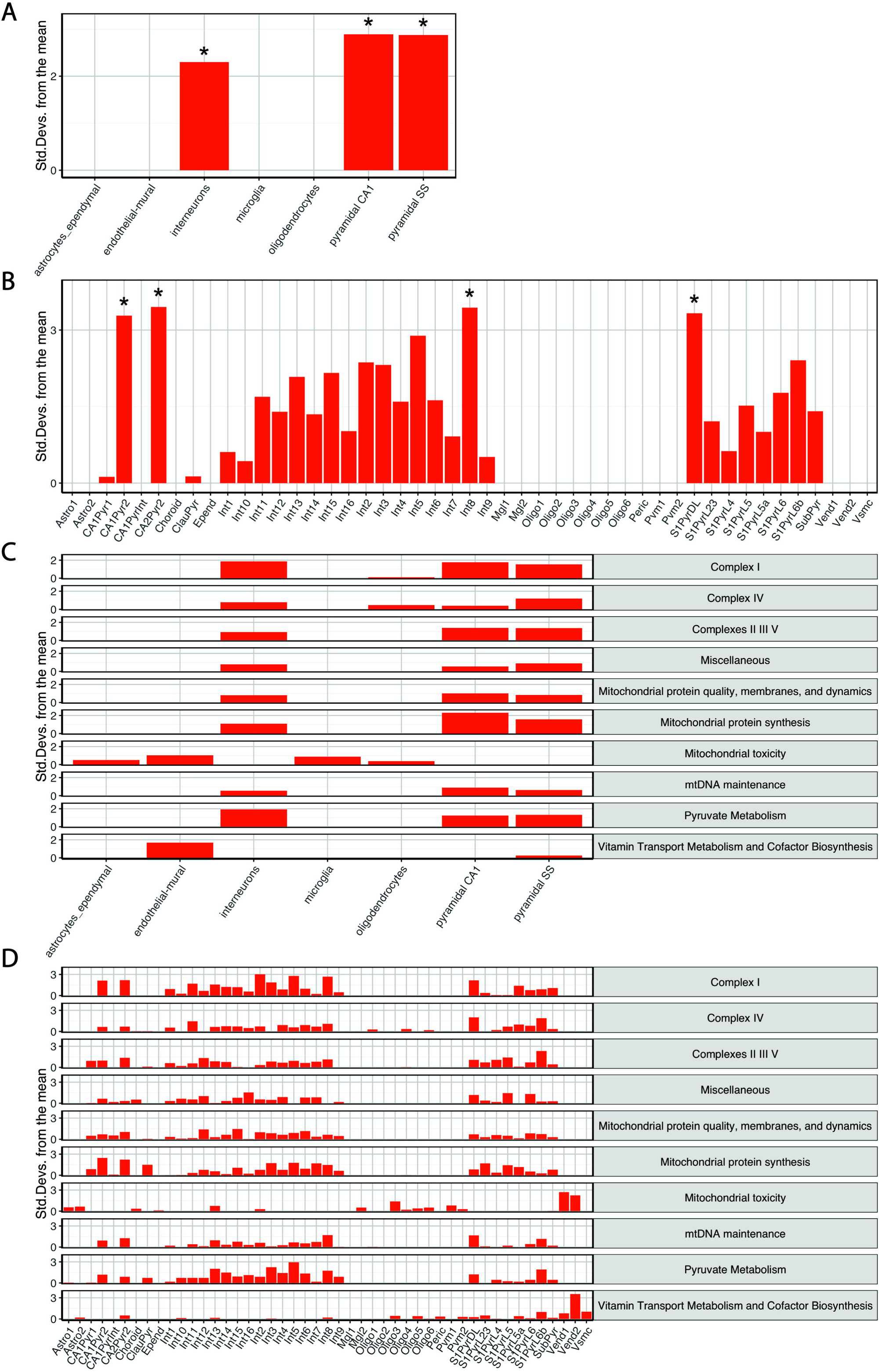
Leigh syndrome associated genes are significantly enriched in interneurons and pyramidal CA1 and SS neurones. Enrichment of Leigh syndrome associated genes in level 1 **(A)** and level 2 **(B)** cell types from the mouse brain single cell RNA-Seq Karolinksa institute superset (reps= 10,000, * denotes significance where Benjamini Hochberg FDR adjusted *P* < 0.05, controlled for GC content and length of genes). EWCE analysis of enrichment of Leigh syndrome associated genes grouped based on gene product function in level 1 **(C)** and level 2 **(D)** cell types from the Karolinska mouse brain superset (reps = 100,000, controlled for GC content and length of genes).

**Figure 3.**
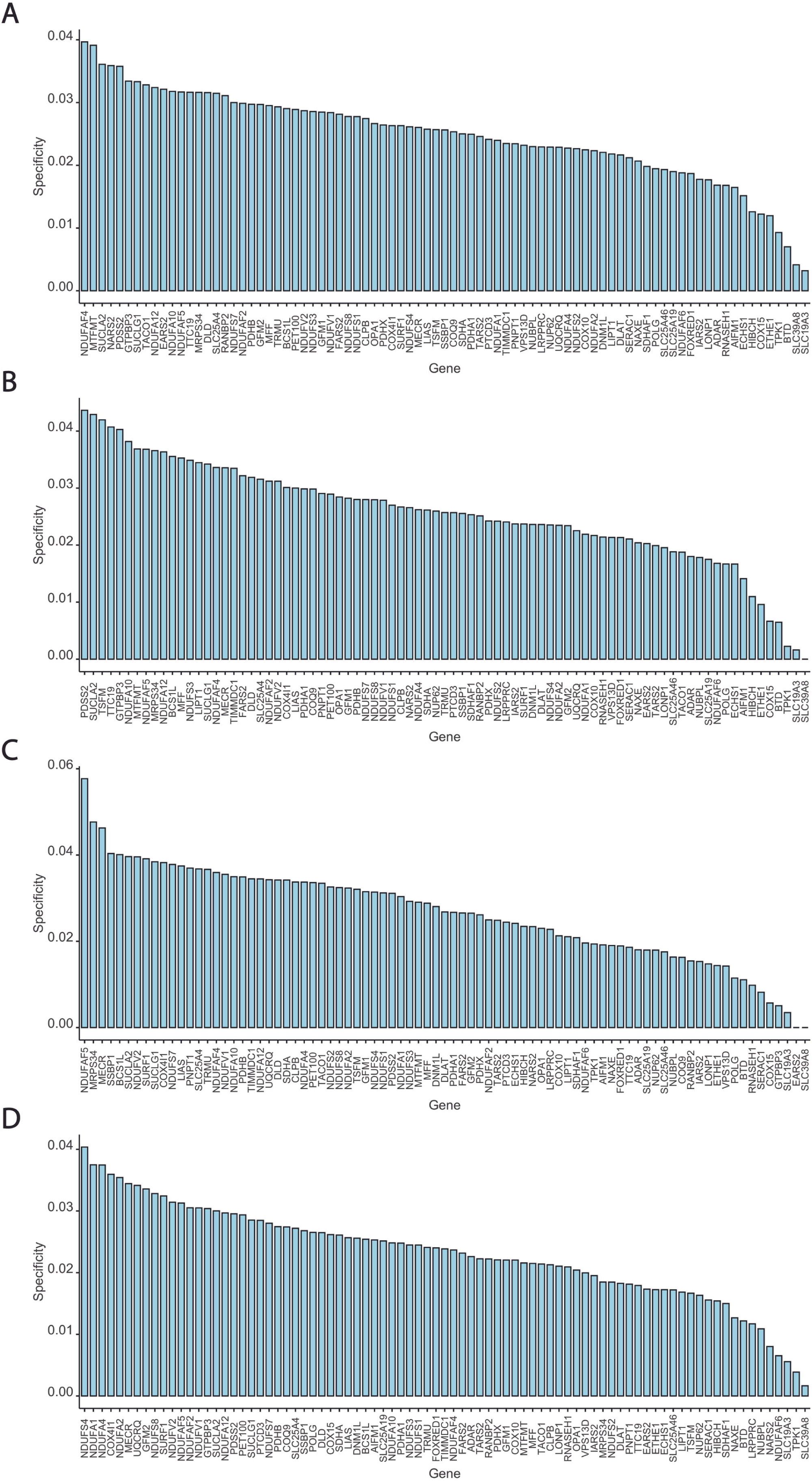
Plot of level 2 cell type specificity values from Karolinska mouse single-cell RNA-Seq superset. Specificity values for Leigh syndrome associated genes within level 2 cell subtypes, CA1 type 2 pyramidal (**A)**, CA2 type 2 pyramidal (**B)**, Interneuron 8 **(C)**, and S1 deep layer pyramidal **(D)** were obtained from Skene et al.^12^ and were generated by dividing the average expression of a gene in a particular cell type by the average expression of the same gene in all other cell types.

## Results

### Leigh syndrome associated genes are highly enriched in the substantia nigra

Utilising transcriptomic data from the UKBEC network within CoExp we sought to identify regions of the brain where Leigh syndrome associated genes were enriched. Of the 113 genes associated with Leigh syndrome (Table 1), 79 were represented within the UKBEC networks. Subsequently, regions of the brain found to have significant enrichment (FDR adjusted *P*-value <0.05) of these genes were identified (Table 3, Fig. 1A). Co-expression network modules with significant enrichment of Leigh syndrome associated genes include four modules of the hippocampus, three modules of the occipital cortex, putamen, substantia nigra, and temporal cortex and two modules of the cerebellum, frontal cortex, medulla, and white matter (Table 2). Modules with enrichment of the most Leigh syndrome associated genes include the red module within the frontal cortex, yellow and light cyan modules of the medulla, the red module for the putamen, grey60 for white matter, and yellow and salmon modules of the substantia nigra (Table 3).

Of the four brain regions with the most Leigh syndrome associated genes identified, namely, substantia nigra, putamen, medulla, and thalamus, the modules with the most Leigh syndrome associated genes were compared to one another to elucidate any common genes (Fig. 1B). Of the genes significantly enriched, nineteen were in common between the four brain regions, namely NDUFS*2, NDUFS4, NDUFA9, NDUFA12, NDUFC2, SDHA, LRPPRC, PDHB, DLAT, DLD, SUCLA2, SUCLG1, NARS2, GFM1, OPA1, SERAC1, MFF, LIAS*, and *RANBP2*. Gene associations unique to the substantia nigra were *NDUFS7, NDUFA10, NDUFA13, EARS2, LONP1*, and *MORC2*. Putaminal-unique genes were *UQCRQ, COX4I1, FBXL4, PNPT1, GFM2, LIPT1, BTD*, and *ETHE1*. Medulla unique genes include *COX8A, NUMPL*, and *TACO1*. Two genes unique to the thalamus are *SLC24A46* and *GTPBP3*. The substantia nigra was found to have enrichment of greater than 64% of the Leigh syndrome associated genes present in the UKBEC network. Of note, nearly 10% of the genes included in the Venny (ie target genes expressed in the substantia nigra, putamen, medulla, or thalamus) are only expressed in the substantia nigra and not the putamen, medulla, or thalamus.

### Leigh syndrome genes are significantly enriched in interneurons and pyramidal cells

To build upon the CoExp findings that Leigh syndrome associated genes are enriched in neuronal tissue, particularly with 64% of genes being enriched within the substantia nigra, we interrogated the Karolinska Institute single cell RNA-Seq mouse brain data set using EWCE^11^. Cells present in level 1 (cell types) of this data set include interneurons, pyramidal cells of the hippocampus (CA1) and somatosensory cortex (SS), astrocytes, glial cells, microglia, oligodendrocytes, astrocytes (ependymal), and endothelial mural cells.

EWCE analysis of the level 1 (higher level cell types) data set revealed that Leigh syndrome associated genes are significantly enriched in interneurons, pyramidal CA1, and pyramidal SS cell types (FDR adjusted *P* = 0.0162, FDR adjusted *P* = 0.0027, FDR adjusted *P* = 0.0036, respectively, Fig. 2A). Notably, OXPHOS enzyme complex subunits and assembly factors were found to be among the top 10 genes in all three cell types where Leigh syndrome associated genes were significantly enriched. EWCE analysis of the cell subtypes (level 2, Fig. 2B) revealed significant enrichment of Leigh associated genes in type 2 pyramidal cells of CA1 (FDR adjusted *P* = 0.0031) and CA2Pyr2 (FDR adjusted *P* = 0.0026). Significant enrichment was also observed in interneuron type 8 (Int8, FDR adjusted *P* = 0.0033) and deep layer pyramidal cells of S1 (S1PyrDL, FDR adjusted *P* = 0.0015).

Further, we sought to determine whether there was differential enrichment of genes based on the function of their products. Following EWCE analysis of the genes subdivided into groups based on similar functions (Table 1), a pattern emerged whereby genes encoding proteins and assembly factors of Complexes I-IV, or involved in mitochondrial protein quality, synthesis, membranes and dynamics were expressed in interneurons and the CA1 and SS pyramidal cells (Fig. 2C). Genes encoding products that are involved in pyruvate metabolism and mtDNA maintenance are similarly expressed. In contrast, genes encoding proteins whose dysfunction results in mitochondrial toxicity were found to be expressed in astrocytes and ependymal cells, endothelial and mural cells, microglia, and oligodendrocytes (Fig. 2C). Leigh syndrome associated genes functioning in vitamin transport metabolism and cofactor biosynthesis were expressed in endothelial and mural cells with some expression in pyramidal SS cells (Fig. 2C). EWCE analysis of the level 2 cell types revealed a similar pattern to that observed for the level 1 data set whereby genes associated with mitochondrial toxicity, vitamin transport metabolism and cofactor biosynthesis had an expression profile different to the rest of the functional groups (Fig. 2D). It should be noted that EWCE findings for the functional groupings did not yield any statistically significant results after correcting for multiple testing when examining both cell types and subtypes, however some trends were observed.

Examining the difference between nuclear and mitochondrial encoded genes associated with Leigh syndrome was inconclusive since none of the mitochondrially encoded genes associated with Leigh syndrome were represented in the data set used for the EWCE analysis. In view of this, we utilised the STRING database^17^ to identify proteins interacting with products from our mitochondrial encoded target gene list and generated an extended ‘mitochondrial’ gene list (Table S3). EWCE analysis of the extended gene list revealed a similar pattern of expression in interneurons, pyramidal CA1, and pyramidal SS cells, however the findings were not statistically significant (Fig. S2). Of note, the STRING-database identified 19 interactors (*MT-CYB, MT-ND4L, MT-ATP8, NDUFA6, NDUFS5, NDUFS6, NDUFAB1, UQCRQ2, COX5A, COX5B, COX6A1, COX6B1, COX6C, COX7A2, COX7C, ATP5A1, ATP5B, ATP5D*, and *ATP5F1*) which are not currently identified as causes of Leigh syndrome but are possible candidate genes for Leigh syndrome.

### Leigh enriched modules are primarily associated with pre-synaptic structures

To understand the role of Leigh syndrome associated genes at the synaptic level and investigate the UKBEC co-expression modules further, we utilised the SynGO database (Fig. 4). CoExp modules containing the largest number of Leigh syndrome associated genes, were found to be significantly enriched in synaptic structures. Of the modules of interest, 4 exhibited enrichment in both pre-and post-synaptic structures, with greater enrichment of pre-synaptic genes. The hippocampus and white matter module of interest demonstrated enrichment in only pre-synaptic structures. We were unable to determine reliably whether the remaining modules with enrichment of Leigh syndrome associated genes exhibited a pre-or post-synaptic enrichment owing to a smaller number of differentially expressed genes in these modules.

**Figure 4.**
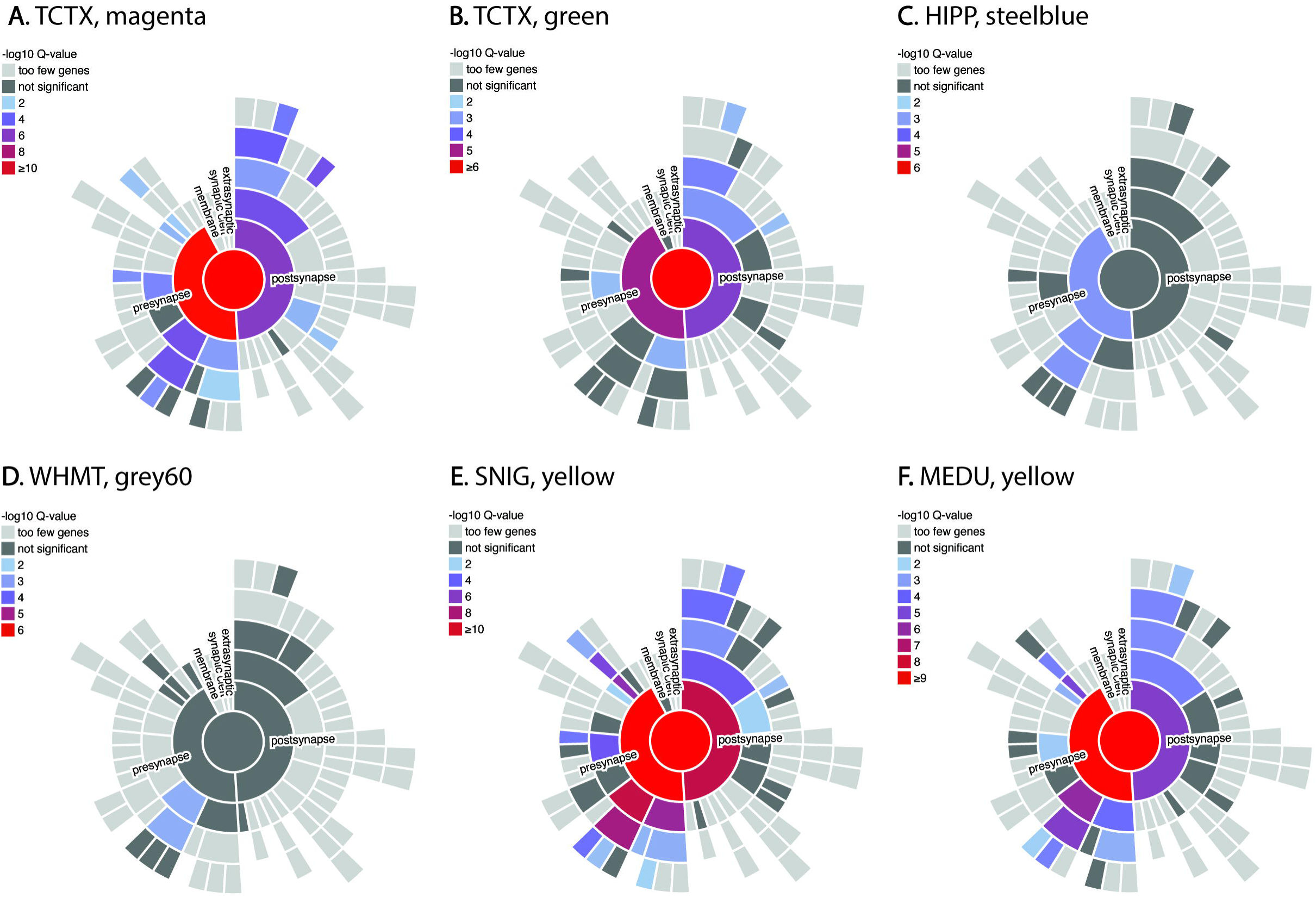
Differential synaptic associations of Leigh syndrome genes in the enriched UKBEC modules, as observed using the SynGO database. This diagram shows that only some of the candidate genes are associated with either presynaptic or postsynaptic structures, or both. It can also be seen that the enriched modules are more strongly linked to presynaptic structures, where mitochondria are required to provide ATP for the secretion and recycling of neurotransmitters. The colour coding is based on enrichment based on q values (-log*P*-values) and concentric rings indicate gene function hierarchy at the synapse with more general terms centrally and specific terms peripherally.

### Co-enrichment of genes associated with Leigh syndrome, epilepsy, Parkinson disease, and schizophrenia

Identification of UKBEC modules in which Leigh syndrome associated genes were enriched allowed us to examine the genetic mechanism of Leigh syndrome compared to other neurological disorders. Owing to some similar clinical features, e.g., dystonia and seizures, we compared Leigh syndrome to Parkinson disease, epilepsy, and schizophrenia (Fig. 5). Modules with significant enrichment of Leigh syndrome associated genes (-logP-value (FDR corrected) > 1.301, or FDR adjusted-*P* value <0.05) were plotted (Fig. 5A). Parkinson disease-causing genes were found to be enriched in substantia nigra yellow (FDR adjusted *P* =0.0271) and white matter grey60 (FDR adjusted *P* =0.049) modules (Fig. 5B). Epilepsy-associated genes are highly enriched within the 10UKBEC medulla yellow (FDR adjusted *P* = 1.4591e-7) and substantia nigra yellow (FDR adjusted *P* = 6.8265e-14) modules (Fig. 5C). Genes involved in schizophrenia were found to be significantly enriched in the 10UKBEC frontal cortex yellow (FDR adjusted *P* =0.0022) and putamen grey60 (FDR adjusted *P* =0.0022) and red (FDR adjusted *P* =0.0482) modules (Fig. 5D).

**Figure 5.**
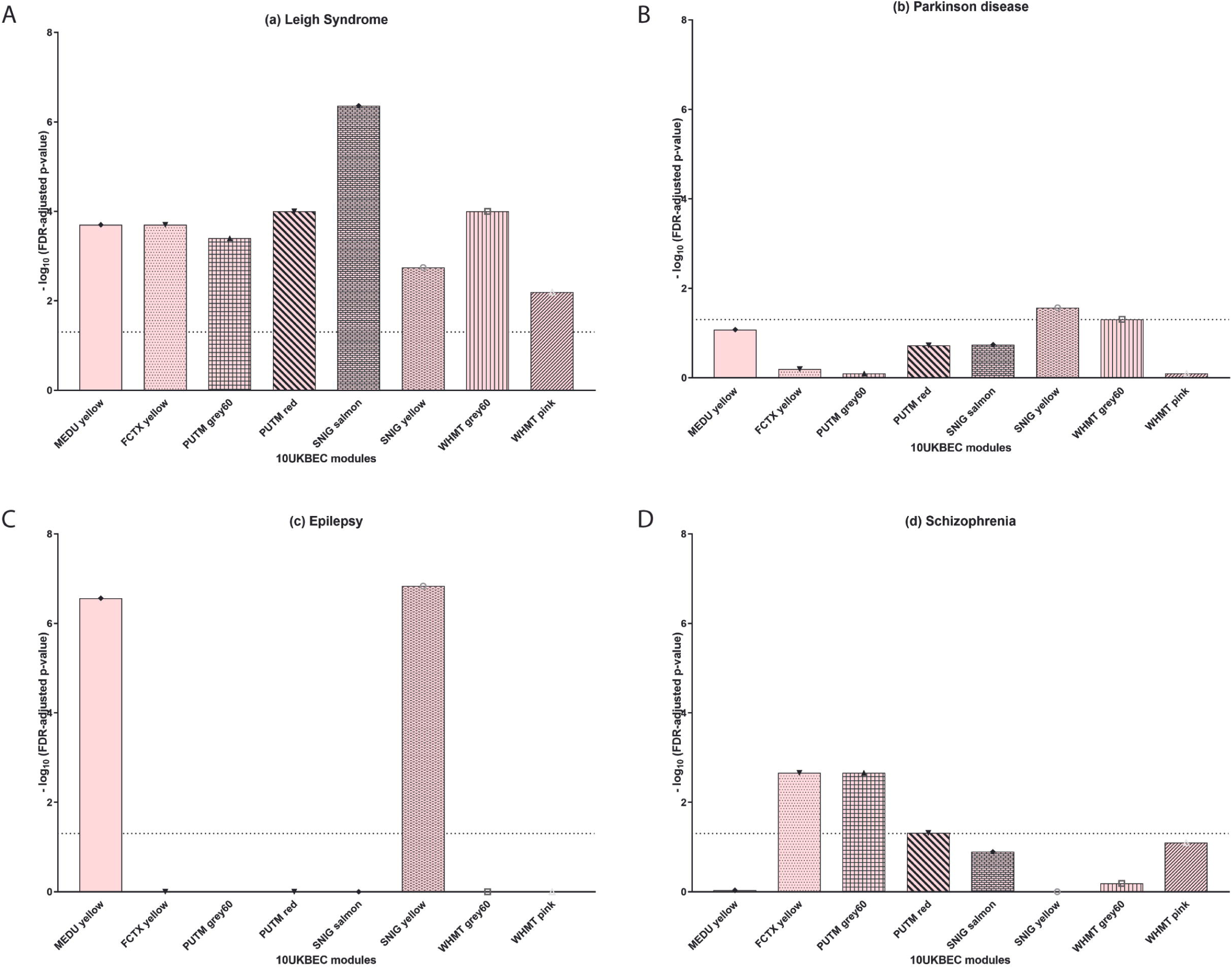
Disease heritability enrichment analysis of Parkinson disease, epilepsy and schizophrenia candidate genes in Leigh syndrome enriched UKBEC modules. The y-axis is the -log10 adjusted *P-* values. The dashed line marks the approximate FDR threshold cut-off for nominal enrichment (approximately 1.301). This figure shows that the medulla ‘yellow’ and substantia nigra ‘yellow’ modules are both enriched for Leigh syndrome **(A)** and epilepsy **(C)** genes, suggesting a possible degree of relatedness in their mechanisms in these two brain regions. The substantia nigra ‘yellow’ module is also enriched for Parkinson disease **(B)** linked genes. Furthermore, frontal cortex yellow, putamen grey60 and red were enriched in schizophrenia **(D)**.

## Discussion

Leigh syndrome is a rare, frequently rapidly progressing neurodegenerative disorder that generally affects children and young infants and often leads to early death. Our current study sought to determine the cell specific expression of Leigh syndrome associated genes in the brain. Here we demonstrate that Leigh syndrome associated genes are enriched in the putamen and substantia nigra. Cell specific analysis of mouse brain transcriptomics revealed significant enrichment in interneurons and hippocampal pyramidal cells. The genes were found to be of relatively similar specificity, with a number of the same genes in the regions with highest specificity, particularly complex I and IV subunits, the complex I assembly factor *NDUFAF5* and genes needed for mtDNA maintenance (especially *SUCLA2*) and mitochondrial gene expression (*GTPBP3, MTFMT* and *MRPS34*). Further, we found that Leigh syndrome associated genes were associated with pre-synaptic structures. Finally, heritability analysis of the co-expression modules revealed a similar enrichment profile in some co-expression modules for epilepsy associated genes with genes associated with Leigh syndrome.

Although Leigh syndrome presents with considerable clinical, biochemical and genetic heterogeneity, the neuropathological features are consistent. Patients exhibit neurodevelopmental delays and progressive decline of the central nervous system with focal bilateral lesions primarily involving the basal ganglia^5^. Lesions in the thalamus, brainstem, spinal cord, and cerebellum may also be present^18^. Of the genes associated with Leigh syndrome represented within the UKBEC network, we found that almost two thirds were enriched within the substantia nigra. This is consistent with clinical presentation of children with Leigh syndrome where necrotising lesions are observed within the basal ganglia, brainstem, and thalamus^19^. Co-expression analyses revealed enrichment of Leigh syndrome associated genes in all ten brain regions within the UKBEC network: cerebellum, frontal cortex, hippocampus, medulla, occipital cortex, putamen, substantia nigra, temporal cortex, thalamus, and white matter. The modules with enrichment of the largest number of genes associated with Leigh syndrome included modules of the medulla, substantia nigra, and putamen. Indeed, pathological lesions in these regions, namely putamen and substantia nigra, are commonly identified on MRI scans in patients with Leigh syndrome and are the most affected areas on neuropathological analysis^18^.

EWCE analysis revealed significant enrichment of Leigh syndrome associated genes in interneurons and hippocampal pyramidal CA1 neurones and SS cells of the somatosensory cortex. Subtype analysis revealed significant enrichment within type 2 pyramidal neurones of CA1 and CA2 regions of the brain, interneuron type 8, and the deep layer pyramidal neurones of the primary somatosensory cortex. The hippocampus possesses a tri-synaptic circuitry involving neurones of the dentate gyrus, CA3 and CA1 areas. CA1 pyramidal cells of the hippocampus represent a key output node of hippocampal memory circuits, with widespread projections^20^. Specifically, type 2 CA1 pyramidal cells have been found to associate with mitochondrial function and have also previously been correlated with the rate of firing and length of projections of cortical neurones ^11,21^. Further, a close relationship between oxygen consumption and gamma wave oscillations within the hippocampus was suggested as there was correlation between spatial expression of complex I subunits and gamma oscillations in mice and rats^21^. These oscillations are thought to function in sensory processing and memory formation^22^. Late onset Leigh syndrome presents with memory loss symptoms, which is consistent with our findings of enrichment of Leigh syndrome associated genes within CA1 and CA2 hippocampal neurones. Loss of mitochondrial function may result in inadequate or loss of fast neuronal network gamma oscillations which occur in hippocampal pyramidal neurones and thus impaired memory formation. The somatosensory cortex receives and processes afferent somatosensory input and contributes to the integration of both somatic and proprioceptive input required for skilled movement. Therefore, mutations in genes enriched in SS pyramidal cells of somatosensory cortex could contribute to the ataxia, dystonia, and movement disorders observed in patients with Leigh syndrome.

We did not find any enrichment in glial cells of Leigh syndrome associated genes in most of the functional groups analysed. Some level of expression within astrocytes and oligodendrocytes was observed in genes whose products are involved in mitochondrial toxicity. Lesions observed in Leigh syndrome are characterised by necrosis, gliosis, demyelination, spongiosis, and capillary proliferation ^19^. The lack of enrichment of Leigh syndrome associated genes in glial cells may be because glycolysis is the main source of energy in astrocytes as opposed to oxidative metabolism in neurones^23,24^. Further, due to impaired mitochondrial functioning and thus neuronal damage and inflammation, reactive gliosis occurs. Morales et al.^25^ have demonstrated impaired corticogenesis in cerebral organoids generated from induced pluripotent stem cells derived from Leigh syndrome patient fibroblasts. Organoids harbouring mutations in *DLD*, PDH, or an *MT-ATP6*/PDH double mutant had increased astrocyte markers levels at both mRNA and protein level. Previous genetic fate mapping has suggested that mature astroglial cells can de-differentiate and resume proliferation^26,27^. Thus, in Leigh syndrome, the upregulation of astrocytes can be due to a reactive process in response to neuronal damage or may be a result of increased cells differentiating down the astroglial lineage.

Functional grouping of Leigh syndrome associated genes revealed that mitochondrial toxicity and vitamin transport metabolism and co-factor biosynthesis demonstrated an expression pattern different to that of all other functional groups. Furthermore, genes associated with mitochondrial toxicity were expressed in both subtypes of astrocytes, those distributed throughout the cortex and those closely associated with the pia^11^. The genes within the mitochondrial toxicity functional grouping include *ETHE1* and *SQOR* involved in sulphide metabolism, *ECHS1* and *HIBCH* associated with valine degradation, and finally *NAXE* and *SLC39A8* with products needed for mitochondrial detoxification. Recent research has indicated that mitochondria are key in regulating astrocyte functions such as, calcium (Ca^2+^) signalling, fatty-acid metabolism, glutamate-glutamine cycling, antioxidant production, and neuroinflammatory activation^28,29^. These observations lead to the hypothesis that in Parkinson disease, mitochondrial dysfunction, through a gain of function mechanism, results in overactive inflammatory function of astrocytes, thereby negatively altering dopaminergic neuronal health.

Enrichment of Leigh syndrome associated genes in the interneurons, pyramidal CA1 and SS cells were found to be similar across most of the genes of interest. It is worth noting that genes encoding OXPHOS enzymes or assembly factors made up half of the genes in the top ten genes in both pyramidal cell types and three of the interneuron cell types, suggesting that Leigh syndrome is related to a primary mitochondrial dysfunction. Furthermore, *SLC19A3* and *SLC39A8* were the least enriched in these three cell types. Interestingly, these two genes encode non-mitochondrial proteins, further suggesting that there is an underlying primary mitochondrial dysfunction in neuronal cells in patients with Leigh syndrome. This could be suggestive of a yet undiscovered mechanism by which *SLC19A3* and *SLC39A8* cause mitochondrial dysfunction or that there is a different pathomechanism by which *SLC19A3* and *SLC39A8* mutations result in a Leigh like syndrome as opposed to Leigh syndrome caused by a primary mitochondrial dysfunction. Given that Leigh syndrome presents on a spectrum with a wide range of disorders and the underlying pathology is due to aberrant mitochondrial function caused by mutations in nuclear or mitochondrial encoded proteins, the finding that no single gene drives the enrichment is as expected. Furthermore, given that the genes encode intracellular proteins and subunits of mitochondrial proteins which are ubiquitously expressed in all human tissues with the exception of mature erythrocytes, such a finding is not unexpected.

Neurones are energy demanding cells that require significant amounts of ATP to generate action potentials and thus normal mitochondrial function is essential for this purpose. Multiple neurodegenerative pathways, including those implicated in Alzheimer^30^, Parkinson^31,32^ and Huntington diseases^33^, are thought to converge on energy failure and thus result in early degeneration of the synaptic terminal, or the pre-synaptic terminal^34^. Canonically, mitochondria are thought to function to provide ATP to maintain the electrochemical gradient and recycling of synaptic vesicles as well as a Ca^2+^ buffer to allow for tight spatial and temporal control of neurotransmitter release^35^. Furthermore, synaptic development and subsequent pruning and plasticity is key for normal functioning of neuronal circuitry. An important factor for adequate development and remodelling of synapses is protein synthesis, another energy demanding process dependent on mitochondrial function^35^. Further, the anti-apoptotic factor BCL-XL (B-cell lymphoma extra-large) has been shown to drive presynaptic maturation via increased activity of dynamin related protein 1 (encoded by *DNM1L*), a protein needed for mitochondrial fission and thereby mitochondrial activity in the presynaptic terminal^36,37^. Both dominant and recessive mutations in *DNM1L* have been linked to Leigh syndrome. Our analyses revealed enrichment of *DNM1L* in the yellow module of the substantia nigra which was found to have significant association with pre-synaptic structures. Previous reports have demonstrated defective bioenergetics^38,39^ in cells derived from patients with Leigh syndrome and more recently, impaired Ca^2+^ homeostasis^40^, illustrating the importance of mitochondria at neuronal synapses. Despite the above, it is well documented that mitochondria are not present at all pre-synaptic terminals^41,42^, and thus it remains unclear whether mitochondria are essential at synapses. Our findings show that certain co-expression modules are significantly associated with pre-synaptic structures while others are not, suggesting that perhaps the expression pattern mirrors that of mitochondrial distribution within neurones. A limitation of this study in determining whether the remaining modules are associated with pre-or post-synaptic structures is the small number of genes associated with these modules.

The inheritance pattern of Leigh syndrome can be either maternally inherited in the case of defective mitochondrial genes, or autosomal recessive, X-linked or dominant if the defects are in nuclear encoded genes. Epilepsy associated genes were found to be enriched in the yellow co-expression modules of the substantia nigra and medulla, modules in which Leigh syndrome genes are also enriched. This correlates with previous studies showing the comorbidity of epilepsy in neurometabolic disorders including Leigh syndrome. A recent systematic review and meta-analysis of 385 patients with Leigh syndrome reported in five studies demonstrated that epilepsy occurs in a third of patients ^43^. Previous literature has also consistently cited epilepsy as one of the clinical features present that may be present in patients with Leigh syndrome, suggesting that there may be a common underlying neuropathological mechanism between epilepsy and Leigh syndrome^44^.

Dystonia, spasticity, ataxia, and tremor are neurological signs that may be present in Leigh syndrome and may also be observed in Parkinson disease. For this reason, we tested whether the genetic basis of Parkinson disease and Leigh syndrome may be linked. Enrichment in the yellow module of the substantia nigra and grey60 module of white matter was observed for Parkinson disease, both regions that were also enriched for Leigh syndrome associated genes. It is well known that Parkinson disease is due to a loss of dopaminergic neurones of the pars compacta of the substantia nigra^45,46^, and many patients with Leigh syndrome have lesions involving the basal ganglia^18^, suggesting there may be a common underlying pathophysiology between the parkinsonism observed in some patients with Leigh syndrome and Parkinson disease. Despite this, no definitive heritability pattern was elucidated, suggesting that the similar clinical features observed may be due to different pathological mechanisms resulting in the same phenotype, or that genes that are not well represented in the studied networks may confer a different inheritance pattern and module enrichment that may reveal further insights. One case with adult onset Leigh syndrome and a mutation in *MTFMT* had atypical parkinsonism^47^. The authors suggested that the dopaminergic neurones in patients with Leigh syndrome may be more susceptible to parkinsonism and Parkinson disease due to impaired OXPHOS leading to extrapyramidal symptoms. Currently, little is known about dopamine transport and signalling in patients with Leigh syndrome and thus future research is warranted to elucidate potential links between Parkinson disease and Leigh syndrome.

Finally, psychiatric symptoms can be present in patients with Leigh syndrome and there is some evidence suggesting that mitochondrial dysfunction is key in the pathophysiology of schizophrenia^48^. For these reasons, we examined the enrichment pattern of schizophrenia associated genes. We found significant enrichment in frontal cortex yellow and putamen grey60 and red for schizophrenia associated genes, modules in which significant enrichment of Leigh syndrome associated genes was also identified. Across the three disorders we compared modules demonstrating enrichment of disease associated genes, it can be noted that the findings for Parkinson disease, epilepsy, and schizophrenia when combined do result in a pattern similar to that observed for Leigh syndrome. This suggests that the underlying pathology of Leigh syndrome may perhaps be predisposing affected patients to develop parkinsonian phenotypes, psychiatric disorders, or seizures.

## Limitations

To our knowledge, this is the first study examining nervous tissue cell specific expression analysis of the genes associated with Leigh syndrome. Despite the novelty of our study, it possesses limitations. Firstly, our study is limited in the availability of high-quality cell specific brain transcriptomic data. The UKBEC co-expression networks utilised in our analyses did not possess data on all of the Leigh syndrome associated genes within all tissues of interest. More specifically none of the 16 mitochondrially encoded genes linked to Leigh syndrome were present within the data set and thus our synaptic enrichment and heritability studies data are incomplete, as the level of enrichment may be an underestimation. For this reason, we were also unable to detect these genes within the specificity analysis. We also acknowledge that the UKBEC data set is an adult human brain data set while Leigh syndrome is primarily paediatric in onset. We are limited in that there are no publicly available paediatric brain data sets and thus perhaps the findings of our study may be underestimated. Secondly, for our EWCE analysis we utilised mouse brain data for its completeness, however once again not all of the Leigh syndrome associated genes were present within the data set, particularly none of the mitochondrial encoded genes. We believe it is for this reason we were unable to identify any significant associations for mitochondrial encoded genes in the functional analysis. We were able to demonstrate enrichment in neuronal cells as opposed to astroglial cells, however, we do appreciate there may be species differences. We were unable to determine whether the spatial expression within brain cells was related to the mode of inheritance of genes associated with Leigh syndrome. As previously mentioned, this was due to poor representation of the mitochondrially encoded genes within the UKBEC network as well as within the mouse single cell RNA-Seq data set used for EWCE. Given this, our findings could potentially underestimate the underlying mitochondrial aetiology of Leigh syndrome. Thirdly, given that Leigh syndrome is genetically heterogeneous with mutations in many different genes encoding very different products having been implicated, it is likely that there are other mutations that cause Leigh syndrome that have yet to be identified. Thus, the findings of our study may change over time as new Leigh syndrome causing gene defects are identified.

## Conclusions

Taken together, our study enabled us to identify specific cells and brain regions that demonstrate enrichment of genes that are associated with Leigh syndrome. We demonstrated that genes associated with the development of Leigh syndrome are expressed within brain regions most affected in patients with Leigh syndrome, namely, putamen, substantia nigra, thalamus, and medulla. Transcriptomic analyses revealed significant enrichment of Leigh syndrome associated genes in neurones, particularly interneurons and pyramidal cells of the hippocampus and somatosensory cortex. More specifically, genes with products involved in mitochondrial toxicity and vitamin transport metabolism and cofactor biosynthesis revealed a different expression pattern with enrichment in astroglial cells as opposed to neurones for the other functional groups. We also show that these genes are preferentially associated with presynaptic structures. Finally, we show that Leigh syndrome associated genes share some enrichment modules with Parkinson disease, epilepsy, and schizophrenia.

## Acknowledgements

We thank Professor Mina Ryten for helpful discussions. All research at Great Ormond Street Hospital NHS Foundation Trust and UCL Great Ormond Street Institute of Child Health is made possible by the NIHR Great Ormond Street Hospital Biomedical Research Centre. The views expressed are those of the author(s) and not necessarily those of the NHS, the NIHR or the Department of Health and Social Care.

## Supplementary material

Supplementary material is available at *Brain* online.

